# Systematic benchmarks reveal DNA polymerase combinations for high-quality genome assembly

**DOI:** 10.64898/2026.11.12.766154

**Authors:** Bernhard Bein, Charlotte Gerheim, Nicola Gscheidel, Lucas Canesin, Alexander Ben Hamadou, Clément Schneider, Tilman Schell, Xueling Yi, Carles Galià-Camps, Julio Hechavarria-Cueria, Juniper A. Lake, Ian McLaughlin, Sylke Winkler, Carola Greve, Michael Hiller

## Abstract

**Background:** A fundamental challenge in biodiversity genomics is generating long-read sequencing data and high-quality genome assemblies from samples lacking gold-standard flash-frozen tissue, including ethanol-preserved specimens, minimally invasive biopsies, and small-bodied organisms. Amplification-based library preparation enables long-read sequencing of such samples, but assembly quality critically depends on the polymerase used for amplification.

**Results:** We present a systematic benchmark of eleven long-range DNA polymerases for PacBio HiFi sequencing and genome assembly across three taxonomically diverse species: a springtail, a sea slug, and a bat. We show that polymerase choice substantially affects assembly contiguity and gene completeness, primarily caused by polymerase-intrinsic GC-dependent amplification bias and read coverage heterogeneity. Five polymerases consistently produced assemblies with comparatively high contiguity and completeness, with some enzymes outperforming the polymerase currently used in the PacBio Ampli-Fi protocol. Importantly, combining reads amplified with different polymerases frequently resulted in superior assemblies, because polymerases complement each other by covering regions where others drop out. Combining reads amplified with different polymerases, we generated haplotype-resolved assemblies of the ∼1 Gb giant springtail and ∼2 Gb bat genome and improved the sea slug reference genome. Using the human GIAB benchmark, we further show that optimized amplification strategies improve variant-calling accuracy, highlighting their utility in medical and clinical research. To facilitate polymerase benchmarking across additional taxa, we provide the polymeval pipeline (https://github.com/casparbein/polymeval), which automates the computational workflow.

**Conclusion:** Our study demonstrates that optimized amplification-based HiFi sequencing generates assemblies with >1 Mb contig N50 and >90% gene completeness, providing a toolkit to produce long-read genome assemblies from challenging samples.

## Background

Large-scale genome sequencing initiatives such as the Vertebrate Genomes Project (VGP), Darwin Tree of Life (DToL), Bird 10,000 Genomes Project (B10K), Bat1K, European Reference Genome Atlas (ERGA), Global Invertebrate Genomics Alliance (GIGA), and related efforts have transformed genomics by generating high-quality reference genomes across the tree of life [1–7]. These initiatives contribute to the goals of the Earth BioGenome Project, which aims to sequence and characterize the genomes of all eukaryotic biodiversity [8].

Long-read (PacBio HiFi, Oxford Nanopore) sequencing and long-range chromatin contact information (Hi-C) have become the state-of-the-art for *de novo* genome assembly, routinely producing highly contiguous, near-complete chromosome-level assemblies, even for complex and repeat-rich genomes [1,4,9,10]. Building on these advances, complete telomere-to-telomere (T2T) assemblies have now been generated for multiple species [10–14]. These high-quality assemblies substantially improve gene annotation and structural variant detection, providing a more complete picture of genome architecture [15,16]. Such genomes also provide a powerful basis for comparative and evolutionary analyses, helping to resolve phylogenetic relationships and uncover principles of genome evolution [17,18]. Furthermore, they enable a better characterization of genetic diversity, thereby supporting species conservation efforts [19].

Despite these advances, obtaining suitable biological samples remains a major bottleneck for long-read sequencing. Long-read library construction typically requires sufficient amounts of high-molecular-weight (HMW) DNA, ideally extracted from flash-frozen tissue samples. Such material is often unavailable or difficult to obtain from endangered or protected species, organisms inhabiting remote environments, or situations in which only minimally-invasive sampling (e.g. biopsies) is possible. In addition, a substantial fraction of eukaryotic biodiversity consists of very small organisms, such as arthropods, gastrotrichs, tardigrades, and nematodes, that provide extremely limited amounts of DNA per individual [20–22]. It has been estimated that for about half of all eukaryotic species less than 1 ng of HMW DNA can be extracted from a single specimen [8]. Adding to these challenges, co-extracted substances such as mucus or naturally occurring metabolites can inhibit long-read sequencing [23,24], and DNA damage introduced during sampling or preservation can further reduce sequencing output.

To overcome these challenges, amplification of genomic DNA prior to sequencing library preparation has emerged as a promising strategy. Amplification-based approaches, such as PImS [25] and the PacBio Ultra-Low Input (ULI) protocol, enable the sequencing of tiny specimens and have been successfully applied to hundreds of species by the Darwin Tree of Life consortium [26]. The ULI protocol has also been utilized for detecting medically relevant variants from limited human samples, such as tissue biopsies and blood samples from patients or newborns [27,28].

The main drawback of whole-genome, long-range PCR DNA amplification is amplification bias. Preferential amplification of specific genomic regions, often influenced by GC-content or noncanonical DNA structures, leads to uneven read coverage and compromises *de novo* genome assembly [29–31]. We recently demonstrated that the choice of the DNA polymerase critically influences these biases. Specifically, replacing or complementing the two long-range polymerases used in the original ULI protocol with the KOD Xtreme Hot Start DNA polymerase substantially reduced amplification bias across taxonomically diverse genomes, including human, sloth, mollusks, and an arthropod, leading to substantially increased assembly contiguity and completeness [24]. Following this work, KODXtreme was adopted in PacBio’s new amplification-based “Ampli-Fi” library preparation protocol [32] and was also shown to successfully amplify formalin-treated, previously cross-linked DNA as part of the CiFi protocol [33]. However, KODXtreme is only one of several commercially available long-range PCR enzymes, many of which have not been systematically evaluated for their performance in whole-genome / long-range PCR amplification, long-read sequencing, or de novo assembly.

Here, we present a comprehensive benchmark of eleven commercially available long-range polymerases, assessing their amplification bias, genome coverage, and impact on assembly quality across three taxonomically diverse species. We show that polymerases differ substantially in the quality of the assemblies they produce, driven primarily by GC-dependent amplification bias and read coverage heterogeneity. Importantly, combinations of the top-performing polymerases frequently generate superior assemblies, collectively compensating for coverage dropouts of individual enzymes. Our results establish a best-practice framework for obtaining high-quality genomes from limited or challenging samples, facilitating reference genome generation across eukaryotic diversity.

## Results

### Amplifying and sequencing DNA with eleven different polymerases

To investigate the performance of different polymerases to amplify DNA for HiFi sequencing and genome assembly, we initially selected a panel of ten polymerases, advertised by manufacturers as capable of amplifying kilobase-long DNA fragments (Table 1). These polymerases have been used in previous studies (Table S1), but a side-by-side comparison is lacking. We systematically tested their performance on three taxonomically diverse species, namely a mollusk (the sea slug *Elysia timida*), an arthropod (the giant springtail *Tetrodontophora bielanensis*) and a mammal (Seba’s short-tailed bat, *Carollia perspicillata*), hereafter referred to by their genus names (*Elysia*, *Tetrodontophora*, and *Carollia*). These species were chosen because they present different challenges for genome sequencing: secondary metabolites inhibit sequencing, a known issue in mollusks [23], organisms with small body sizes such as springtails provide limited amounts of DNA [20], and for protected species, such as many bats, sample acquisition is difficult.

**Table 1:** Polymerases benchmarked in this study. Abbreviation refers to the names we use in the text and Figures.

| **Abbreviation** | **Polymerase Name** | **Manufacturer** | **Notes** |
|---|---|---|---|
| LongAmpTaq | LongAmp® Taq DNA Polymerase | New England Biolabs |  |
| Equinox | Equinox HC Library Amplification Kit | Watchmaker Genomics |  |
| TaKaRaExPremier | Ex Premier DNA Polymerase | TaKaRa Bio |  |
| BiozymB7 | Biozym B7 Hot-Start High Fidelity DNA Polymerase | Biozym Scientific |  |
| PlatinumSuperFi | Invitrogen Platinum™ SuperFi™ II PCR Master Mix | Thermo Fisher Scientific |  |
| repliQa | repliQa® HiFi ToughMix® | Quantabio |  |
| KODXtreme | KOD Xtreme<sup>TM</sup> Hot Start DNA Polymerase | Sigma-Aldrich | Currently used in PacBio’s ‘Ampli-Fi’ protocol |
| NEBNext | NEBNext® High-Fidelity 2X PCR Master Mix (PacBio Ultra-Low DNA Input A) | New England Biolabs | Used in PacBio’s discontinued ‘Ultra Low Input protocol’ |
| TaKaRaGXL | PrimeSTAR® GXL DNA Polymerase (PacBio Ultra-Low DNA Input B) | TaKaRa Bio | Used in PacBio’s discontinued ‘Ultra Low Input protocol’ |
| DeCodiFi | DeCodi-Fi High Fidelity PCR Kit | Kura Biotech |  |
| TaKaRaLongSeq | PrimeSTAR® LongSeq DNA Polymerase | TaKaRa Bio | Added during the study |

To eliminate DNA extraction as a confounding factor, a single DNA extraction was performed from one individual for each of the three species (Figure 1A). The springtail *Tetrodontophora* was deliberately selected due to its comparatively large body size (7–9 mm), which is greater than that of most species in this group, as this enabled us to extract sufficient DNA from a single individual. For each species, long-range PCR amplifications were performed using ten polymerases in combination with a modified PacBio Ampli-Fi SMRTbell library preparation protocol (Figure S1). The resulting polymerase-specific libraries were barcoded to evaluate their performance, equimolarly pooled, and sequenced on two (*Tetrodontophora*) or four (*Elysia*, *Carollia*) SMRT Cells per species on the PacBio Revio platform.

**Figure 1:**
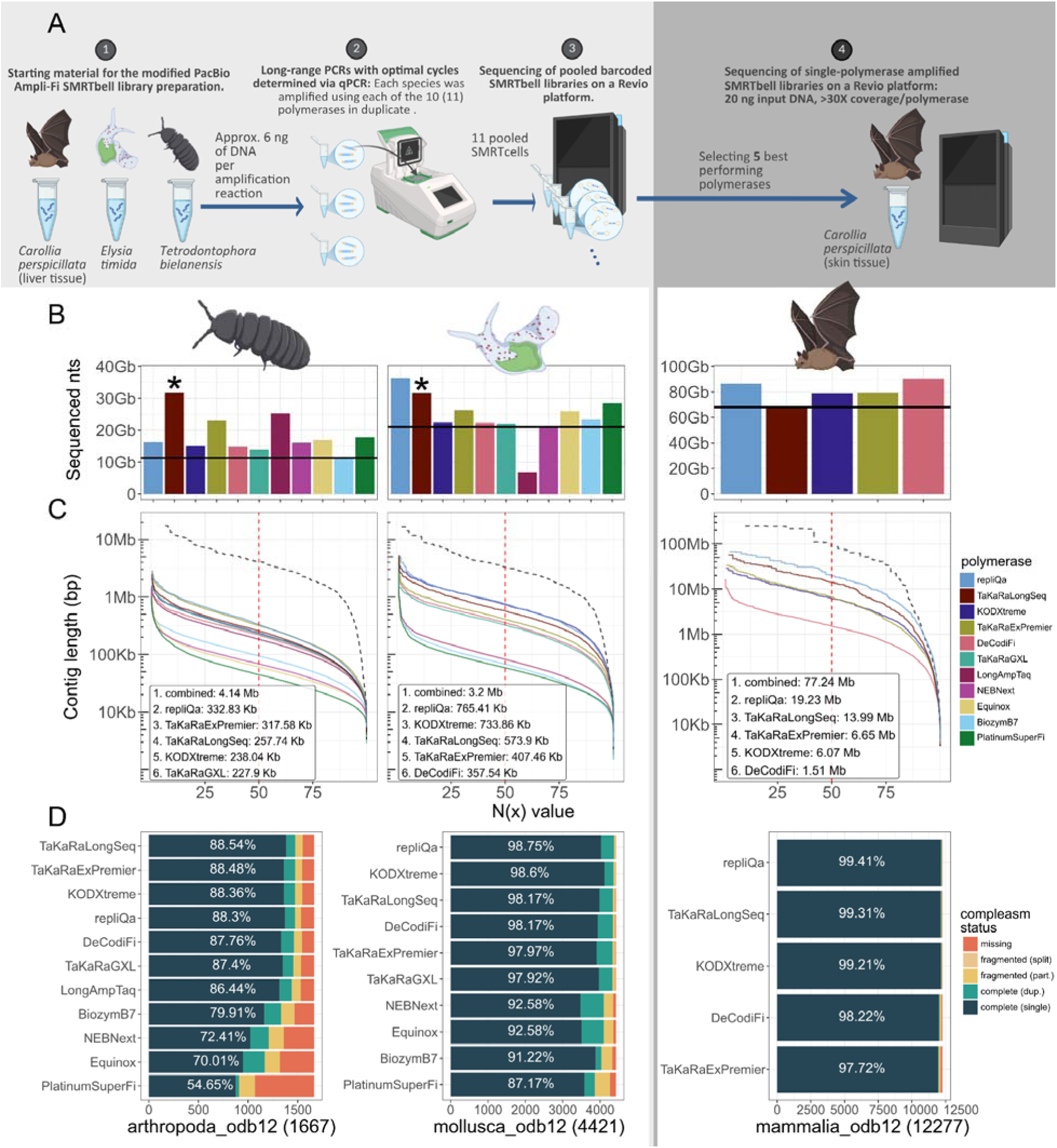
Comparison of polymerase-specific assembly qualities across the three test organisms. (A) Illustration of the workflow, using a single DNA extraction, generating polymerase-specific amplification libraries, and HiFi sequencing (B) Total nucleotide output per polymerase, shown as the total number of nucleotides in HiFi reads. Asterisks indicate that sequencing output for TaKaRaLongSeq is not directly comparable because it received a larger sequencing share. (C) Assembly contiguity from single-polymerase libraries downsampled to equal coverage (different colors) and all libraries combined (dashed line) shown as N(x) plots, where x percent of the assembly (X-axis) consist of contigs of at least N(x) bp (Y-axis). The top five performing polymerases are labeled in the inset in the order of the contig N50. (D) Gene completeness measured by compleasm and sets of near universally-conserved BUSCO odb12 genes of single-polymerase assemblies. Color legend shows the number of genes classified as missing or fragmented in the assembly and genes that are completely present as a single copy or duplicated. (B–D) For *Carollia*, only the deeper-sequenced benchmark of the five best-performing polymerases is shown (all initial polymerases are shown in Figures S3 and S5).

During the course of this study, we became aware of TaKaRaLongSeq, a newly available long-range DNA polymerase (Table 1), and included it as the eleventh polymerase in our benchmarks. For the springtail and sea slug, we generated TaKaRaLongSeq-based libraries using DNA from the same extractions used also for the initial ten polymerases and pooled these libraries on one additional SMRT Cell.

For *Carollia*, we adjusted the experimental design for two reasons. First, no DNA extract remained from the initial experiment, precluding the preparation of a TaKaRaLongSeq-based library from the same extraction. Second, given the larger *Carollia* genome (>2 Gb), the data obtained from the four SMRT Cells in the initial ten-polymerase benchmark provided only ∼8X HiFi read coverage per polymerase. Although this coverage was sufficient to identify polymerases that consistently produced superior assemblies, the resulting assemblies had rather low quality. We therefore performed a second experiment using a single new DNA extraction from the same individual and focused on five polymerases: TaKaRaLongSeq and the four best-performing enzymes identified in the initial benchmark (repliQa, TaKaRaExPremier, KODXtreme, and DeCodiFi; see below). To increase read coverage per polymerase, we sequenced each polymerase-specific library on one SMRT Cell. In the main text, we present the results for these five polymerases for *Carollia*, unless stated otherwise, and provide the results for all ten polymerases from the initial experiment in the Supplementary Information.

### General consistency of polymerase performance across species

For the initial polymerase benchmark, we were able to generate between 6.8 and 36.9 Gb of read data from polymerase-specific amplification libraries (Figures 1B, S2, Table S2); sequencing output for TaKaRaLongSeq is not directly comparable because it was allocated a larger sequencing share. Across the three species, sequencing output was largely consistent for each individual polymerase, with the exception of the LongAmpTaq polymerase, which yielded a high sequencing output for *Tetrodontophora* (25.5 Gb) but the lowest output for *Elysia* (6.8 Gb; Figures 1B, S2). With the exception of LongAmpTaq, which generated very short reads for *Elysia*, read length distributions obtained with the eleven polymerases were broadly similar across the three organisms with median read lengths ranging from 4,127 to 7,322 bp (Figure S2). This indicates that polymerase-specific differences in amplified fragment lengths are largely species independent. Importantly, the fraction of PCR duplicates was generally low in all sequenced libraries (range 0.53 – 4.78%, Table S3), showing that libraries sequenced with long-range PCR are sufficiently complex for genome assembly.

### Substantial differences in polymerase-specific assembly quality

We next evaluated the quality of genome assemblies using read data generated by polymerase-specific amplification. To this end, we removed PCR duplicates and generated contig-level assemblies with hifiasm v0.25.0 [34]. Since assembly quality will depend on the amount of read data, which varies between the polymerases, we removed this confounding factor by downsampling the read data of each polymerase to match the lowest number of bases sequenced across all polymerases for that species (Figures 1B, S3A). Hence, in all following tests, assemblies were generated from an equal number of bases. To evaluate assembly contiguity and completeness, we compared the contig N50 value and the percent of completely detected BUSCO ODB12 genes inferred with compleasm v0.2.9 [35]. These two key metrics should be interpreted as relative among the tested polymerases and are intended to infer the best-performing ones.

For the springtail *Tetrodontophora*, we found that assemblies generated from reads amplified with repliQa, TaKaRaLongSeq, TaKaRaExPremier, KODXtreme, DeCodiFi, TaKaRaGXL, and LongAmpTaq exhibited higher contiguity (contig N50 184-332 kb) and higher gene completeness (arthropoda_odb12 86.4-88.5%) than assemblies generated with the remaining four polymerases (Figure 1C-D). Surprisingly, the assembly sizes were much larger (681 Mb - 1.01 Gb, Figure S4A-B) than expected from available assemblies of related springtails; however, *Tetrodontophora* has never been sequenced. To verify that the large assembly size reflects the actual genome size, we used flow cytometry on a male and a female individual, obtaining genome size estimates of 933 -1099 Mb (Table S4). This indicates that *Tetrodontophora* has the largest known collembolan genome [36] and shows that our assembly size is not artificially inflated. Given the unexpectedly large genome, the estimated coverage after downsampling is only ∼11X, which explains the comparatively low contiguity and completeness; however, our comparisons are relative. Without downsampling, contig N50 values of 1.57 Mb and gene completeness values of 91.66% were achieved (Figure S5, Table S5), comparable to current long-read springtail assemblies [20,37].

For the mollusk *Elysia*, we excluded LongAmpTaq in our downsampling tests, as this polymerase-specific amplification library produced the shortest read lengths and an exceptionally low sequencing output. Instead, we subsampled to match the output of the second-lowest polymerase (NEBNext). Similar to the springtail, we found that assemblies generated from reads amplified with repliQa, TaKaRaExPremier, KODXtreme, DeCodiFi, and TaKaRaLongSeq showed higher contiguity (contig N50 357 - 765 kb) and greater gene completeness (mollusca_odb12, 97.97-98.75%; Figure 1C-D) than assemblies generated with the other five polymerases. Assembly sizes ranged from 626 - 767 Mb and are consistent with 754 Mb size of the available reference genome (Figure S4C-D) [23]. Given the reference genome size, we expect an average coverage of ∼28X per polymerase after downsampling. Assemblies computed without downsampling had contig N50 values of up to 1.32 Mb and gene completeness of up to 98.94% (Figure S5, Table S5).

For the bat *Carollia*, the benchmark of the initial ten polymerases, downsampled to a low read coverage of 8.5X, was broadly consistent with our results for *Elysia and Tetrodontophora* (Figures S3, S5, Table S5). In the subsequent, deeper-sequenced benchmark of the five top-performing polymerases (including TaKaRaLongSeq), repliQa produced the most contiguous assembly (contig N50 19.23 Mb), and repliQa, TaKaRaLongSeq, and KODXtreme yielded the highest gene completeness (mammalia_odb12 99.2-99.4%) (Figure 1C-D, Table S6).

In addition to assembly completeness and contiguity, we also assessed the number of base errors caused by amplification. To this end, we used hifieval [38] to compute the proportion of all sequenced bases corrected by hifiasm’s internal error correction module as a proxy of erroneously sequenced bases. We found that all polymerases except LongAmpTaq had a similarly low percent of corrected bases, averaging 0.01% for *Tetrodontophora* and *Carollia*, and 0.013% for *Elysia* (Figures S2C, S5G-I, S9D, Table S5, S7). LongAmpTaq showed elevated percentages (0.015-0.02%), indicative of a higher number of base errors, in line with results reported by [31].

In summary, five polymerases (repliQa, TaKaRaLongSeq, TaKaRaExPremier, KODXtreme, and DeCodiFi) consistently outperformed the others across three taxonomically diverse species, yielding assemblies with higher contiguity and gene completeness. Pronounced species-specific differences were observed for LongAmpTaq and TaKaRaGXL. LongAmpTaq performed comparatively well for *Tetrodontophora* and *Carollia*, but ranked worst for *Elysia*. TaKaRaGXL showed markedly reduced gene completeness in *Carollia* (Figure S3C), whereas its performance was comparable to other polymerases in *Elysia* and *Tetrodontophora*. Overall, with two notable exceptions, polymerase performance is largely consistent across species, indicating that intrinsic characteristics of these enzymes, rather than sample-specific factors, are the primary determinants of assembly quality.

### Coverage heterogeneity and GC-dependent amplification bias explain assembly quality differences

To investigate the reasons underlying these polymerase-specific sequencing characteristics, we first analyzed the frequency of chimeric reads, which connect non-adjacent genomic regions and hinder assembly.

This analysis requires a reference genome of the three species to which deduplicated amplified reads can be mapped to. Whereas reference genomes are available for *Elysia* and *Carollia* [23,39], *Tetrodontophora* has never been sequenced before. We therefore generated a new haplotype-resolved, chromosome-level reference genome for this species. We pooled HiFi data obtained from the different polymerases (totaling 172 Gb, ∼172X coverage) to generate contigs, and sequenced Arima Hi-C chromosome conformation capture data (∼31X coverage) for scaffolding. The resulting assemblies have contig/scaffold N50 values of 1.46/111.2 Mb and 1.25/104.5 Mb for haplotypes 1 and 2, respectively (Figure S6). For the chimeric read analysis, we used haplotype 1.

The frequency of chimeric sequencing reads was inferred by counting the proportion of supplementary alignments compared to the number of primary alignments for each of the three test species. Consistent with previous observations [24], we found that all long-range PCR amplified libraries exhibited low chimera frequencies, with fewer than 10% of reads having a supplementary alignment, independent of the used polymerase (Figure S7). This suggests that chimeras are not a main reason behind assembly quality differences. In contrast, multiple displacement amplification has been reported to produce on average 2.7 alignments per read, with over 70% of reads producing at least one supplementary alignment [40,41].

A second reason underlying assembly quality differences could be polymerase-specific amplification biases, resulting in uneven read coverage or dropouts in GC-rich or GC-poor genomic regions [30,42]. To quantify read coverage heterogeneity for each polymerase, we tiled the genome into non-overlapping 100 kb windows and computed the interquartile range (IQR), a measure of data spread defined as the difference between the 25th and 75th percentiles, of normalized coverage. We then defined coverage heterogeneity as the median IQR per sample. Furthermore, we computed the low coverage index (LCI), defined as the proportion of the genome with a normalized coverage below 0.5. Both of these metrics differ markedly between the polymerases and correlate with the contiguity and completeness of the assemblies. In particular, the five best-performing polymerases consistently show lower coverage heterogeneity and LCI values than the other polymerases, indicating more even read coverages and less amplification bias (Figures 2A-B, S8A-B).

**Figure 2:**
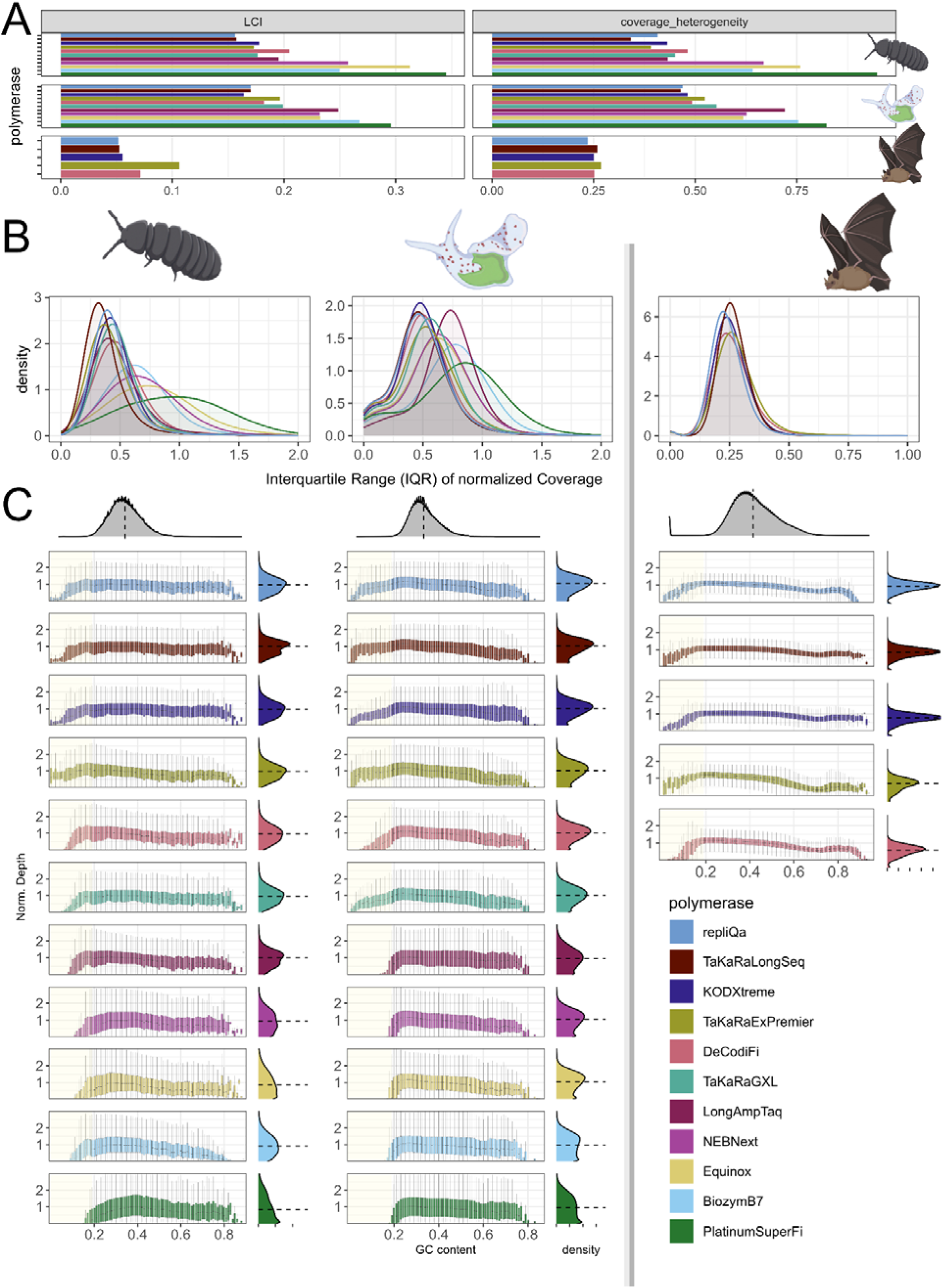
Coverage heterogeneity and GC-dependent dropouts of tested polymerases. (A) Per-polymerase low-coverage index (LCI) and coverage heterogeneity for each of the three test species. LCI is the proportion of the reference genome with less than half of the normalized read coverage. Coverage heterogeneity was quantified as the median interquartile range (IQR) of normalized read coverage across 100kb nonoverlapping windows of the reference genome. (B) Density plots of IQR values calculated in 100 kb nonoverlapping windows of the reference genome. Right-shifted curves indicate more uneven read coverage. (C) Normalized read coverage in 200 bp nonoverlapping windows, partitioned by GC-content in each reference genome window and plotted for each polymerase separately. Boxplots show the 25%, median and 75% quantiles; whiskers extend to 1.5-times the interquartile range (outliers are not depicted). The top row shows the genome-wide GC-content density. Overall normalized coverage densities are shown on the right side. Faint yellow background highlights GC regions where at least one polymerase readset does not have any coverage. Grey lines separate initial benchmarks (left) and deeper-sequenced top-performing polymerases in *Carollia* (right). See Figure S8 for initial benchmarks of *Carollia*.

We next investigated whether GC content accounts for polymerase-specific amplification bias (Figure 2C, Figure S8C). Indeed, polymerases with elevated read coverage variability showed severe dropouts in regions with low GC content (NEBNext, Equinox, PlatinumSuperFi, LongAmpTaq), high GC content (TaKaRaGXL) or both (BiozymB7). In contrast, although the best-performing polymerases also showed decreases in normalized coverage in high- or low-GC regions, they did not experience severe dropouts. In summary, polymerase-specific dropouts in low- or high-GC regions inevitably disrupt contig assembly and are therefore a main contributor to decreased assembly quality.

### Polymerase combinations yield superior assemblies

Building on our previous finding that combining reads from different polymerase amplifications can improve assembly contiguity by compensating for polymerase-specific coverage gaps in genomic regions that individual polymerases fail to amplify [24], we systematically evaluated whether this strategy improves assembly quality across our three test species. To enable a fair comparison, read data from each polymerase was downsampled such that all assemblies were generated from an equal number of sequenced bases.

We first tested all pairwise polymerase combinations. We found that pairs of polymerases that performed poorly in our individual tests (NEBNext, Equinox, PlatinumSuperFi, BiozymB7) never produced top-ranked assemblies in terms of contiguity and gene completeness (Table S7, indicating that they do not complement each other. Combinations involving LongAmpTaq produced comparatively good assemblies for *Tetrodontophora* and *Carollia*, but insufficient coverage prevented its use for *Elysia*. This polymerase also exhibited the highest base error rate across all test species (Figures S2C, S5G–I, Table S5). Combinations involving TaKaRaGXL yielded assemblies with substantially reduced gene completeness in *Carollia*, likely because of pronounced high-GC dropouts (Figure S8C). Given these limitations, LongAmpTaq and TaKaRaGXL were also excluded from subsequent analyses. We therefore focused on the five top-performing polymerases: repliQa, TaKaRaLongSeq, TaKaRaExPremier, KODXtreme, and DeCodiFi. This enabled us to systematically test combinations of three, four and all five (Figure 3A-B), which would have been computationally prohibitive with all eleven polymerases.

**Figure 3:**
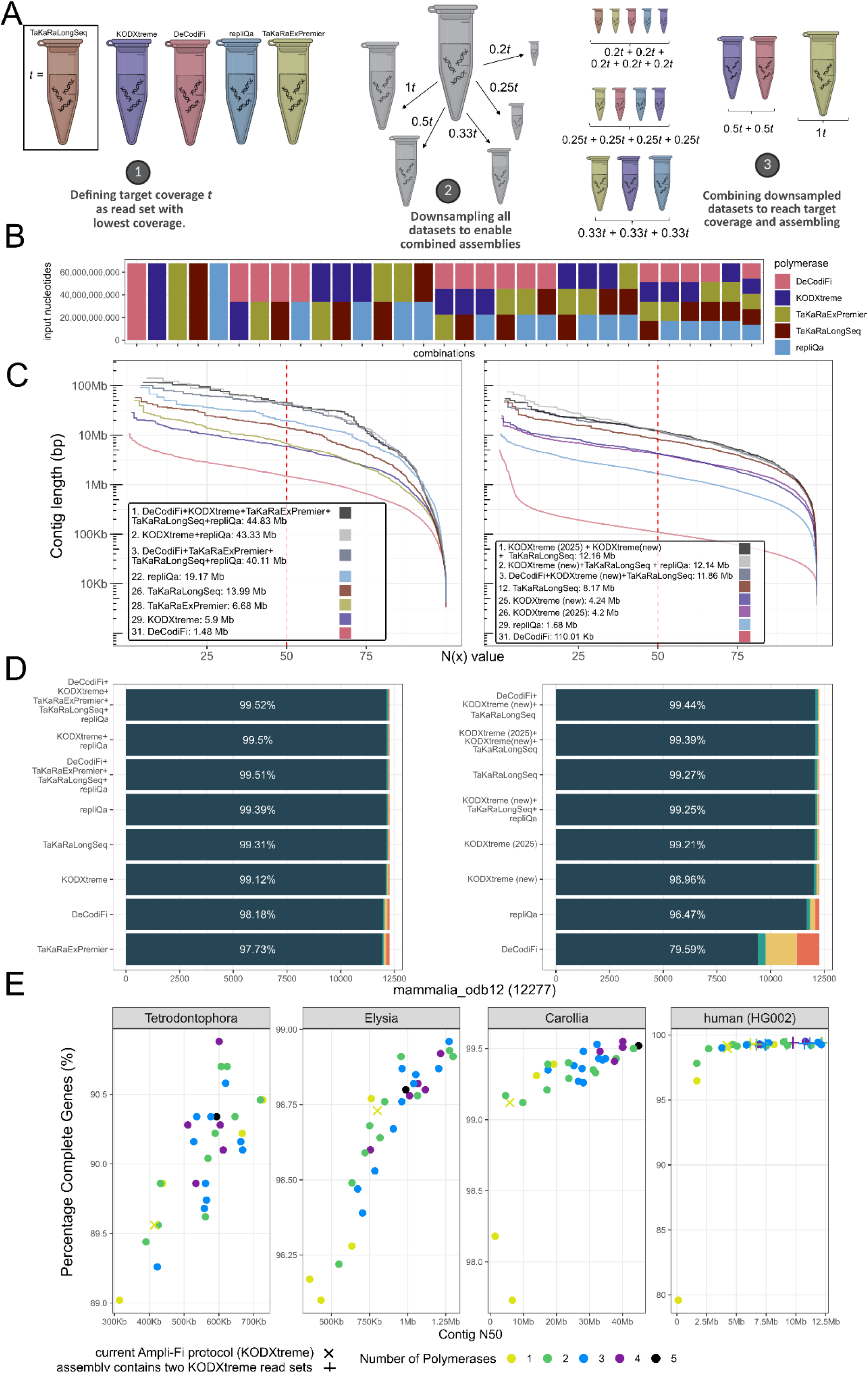
Polymerase read dataset combinations yield more contiguous and complete assemblies. (A) Schematic overview of downsampling and combining of read datasets for assembly for *Carollia*. The dataset with the fewest sequenced nucleotides is chosen as the target coverage t (1). All datasets are then downsampled to 1×t, 0.5×t, 0.33×t, 0.25×t, and 0.2×t (2). Downsampled datasets are combined to always obtain t (3). (B) Overview of downsampled datasets for *Carollia*, equally combining amplified data from one to five polymerases such that all combinations comprise 68 Gb. (C) Assembly N(x) graphs for *Carollia* (left) and human HG002 (right) show that combined-polymerase assemblies outperform single-polymerase assemblies. The best three assemblies (black, light grey, and dark grey) and for comparison the single-polymerase assemblies are indicated in color and their contig N50 is given in the inset. (D) Gene completeness measured by compleasm and sets of near universally-conserved BUSCO odb12 genes of single- and combined-polymerase assemblies. Color legend shows genes that are classified as missing or fragmented in the assembly and genes that are completely present as a single copy or duplicated. The percentage of complete genes is shown in white font. (E) Scatter plot showing contig N50 (X-axis) and gene completeness (Y-axis) of single- and combined polymerase assemblies. Colors indicate the number of different polymerase datasets combined for assembly. Single-polymerase assemblies generated with the Ampli-Fi protocol KODXtreme polymerase are denoted as ‘x’. In the HG002 plot, ‘+’ indicates assemblies where two KODXtreme datasets were included.

Interestingly, for two of the three test species, assemblies generated from polymerase combinations were superior to those produced from individual polymerase data (Table S8, Table S9). While for *Tetrodontophora*, the assembly based solely on repliQa reads outperformed all other combinations in contiguity (contig N50 726 kb, gene completeness 90.46%), for *Elysia* the two polymerase combination DeCodiFi–repliQa increased contiguity from 802 kb to 1.29 Mb and gene completeness from 98.73 to 98.91% relative to the best assembly from single polymerase data (KODXtreme), and the three polymerase combination DeCodiFi– repliQa–TaKaRaLongSeq even yielded a slightly higher gene completeness of 98.96%. For high sequencing-depth *Carollia* data, combining all five polymerases increased contiguity from 19.22 Mb to 44.83 Mb and gene completeness from 99.39% to 99.52% compared with the best single-polymerase assembly (repliQa; Figure 3C-D, Table S9). Furthermore, for *Elysia* and *Carollia*, assemblies generated from three- or four-polymerase combinations were generally better than assemblies generated from pairwise combinations (Figure 3E). This indicates that for some species polymerase-specific coverage dropouts can be compensated by reads from other polymerases.

It is possible that polymerase-specific dropouts are not inherent characteristics of the polymerases themselves, but instead are caused by limited library complexity or random batch effects during DNA extraction, library preparation, amplification, or sequencing. To investigate this, we reanalyzed our two independent *Carollia* read sets and found that only combining reads from different polymerases from either experiment read set consistently improved assembly quality (Tables S10, S11). This indicates that combining reads from different polymerases is more effective at improving assembly quality than repeated amplifications with the same polymerase.

In summary, these tests show that combinations of polymerases produce superior assemblies for some species.

### Polymerases complement each other to recover dropouts

To explore more deeply the reasons contributing to polymerase complementarity, we reasoned that dropouts of individual polymerases, which cause contig breaks, may be compensated by data from other polymerases in a combined assembly. To investigate this, we mapped contigs from polymerase-specific assemblies to the *Carollia* reference genome (Figure 4A), allowing us to identify assembly gaps as genomic regions between adjacent contigs. We then asked (i) whether assembly gaps coincide with decreased read coverage and dropouts of the respective polymerase, and (ii) whether these gaps are covered by reads from other polymerases. Indeed, assembly gaps for all five polymerases generally corresponded to pronounced drops in coverage, often reaching complete dropout, whereas reads from one or more other polymerases cover these gaps (Figure 4B).

**Figure 4:**
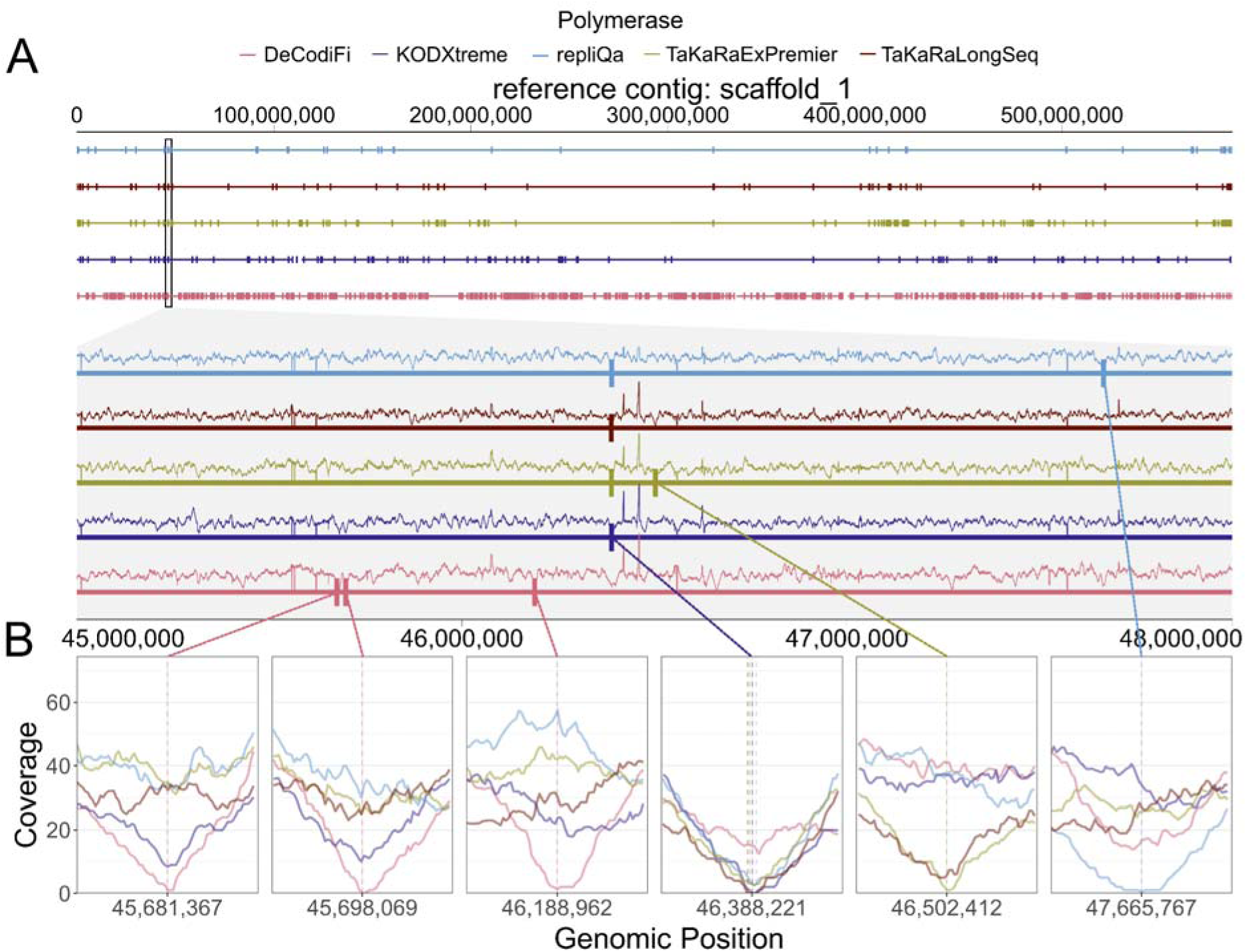
Complementary dropouts of the best five polymerases in *Carollia*. (A) Illustration of the longest scaffold of the reference *Carollia* assembly, showing assembly gaps in single-polymerase assemblies as tickmarks. The grey box zooms into a representative region (scaffold_1 coordinates 45,000,000-48,000,000), showing assembly gaps as boxes and read coverage as lines. (B) Six examples of assembly gaps in single-polymerase assemblies show that these gaps typically coincide with coverage dropouts of the respective polymerase-amplified reads. Colors represent the five different polymerases

Interestingly, we observed non-random, shared read coverage decreases for certain polymerases. For example, regions that are assembly gaps in the DeCodiFi-based assembly typically have reduced KODXtreme read coverage, and vice versa (Figure 5A). This is supported by DeCodiFi- and KODXtreme-based assemblies having the highest number of overlapping assembly gaps (327) and KODXtreme+DeCodiFi assemblies having the lowest contiguity of all pairs (Table S11), suggesting that these two polymerases share similar intrinsic biases. Likewise, TaKaRaLongSeq and TaKaRaExPremier-based assemblies share 87 overlapping assembly gaps and the second lowest contiguity in paired assemblies after DeCodiFi+KODXtreme. Assembly gaps in the DeCodiFi- and KODXtreme-based assemblies consistently showed high coverage from TaKaRaExPremier-amplified reads (Figure 5A), indicating strong complementarity, which is consistent with the high contiguity of DeCodiFi-TaKaRaExPremier pairwise assemblies (Table S11). Assembly gaps specific to TaKaRaExPremier–based assemblies showed only a slight reduction in coverage by the other polymerases, including TaKaRaLongSeq. Finally, assembly gaps in repliQa-based assemblies exhibited a small reduction in coverage by TaKaRaExPremier and DeCodiFi, but near normal KODXtreme and TaKaRaLongSeq coverage (Figure 5A), consistent with a high complementarity between repliQa and KODXtreme/TaKaRaLongSeq and the high quality of repliQa-KODXtreme and especially repliQa-TaKaRaLongSeq assemblies (Tables S11).

**Figure 5:**
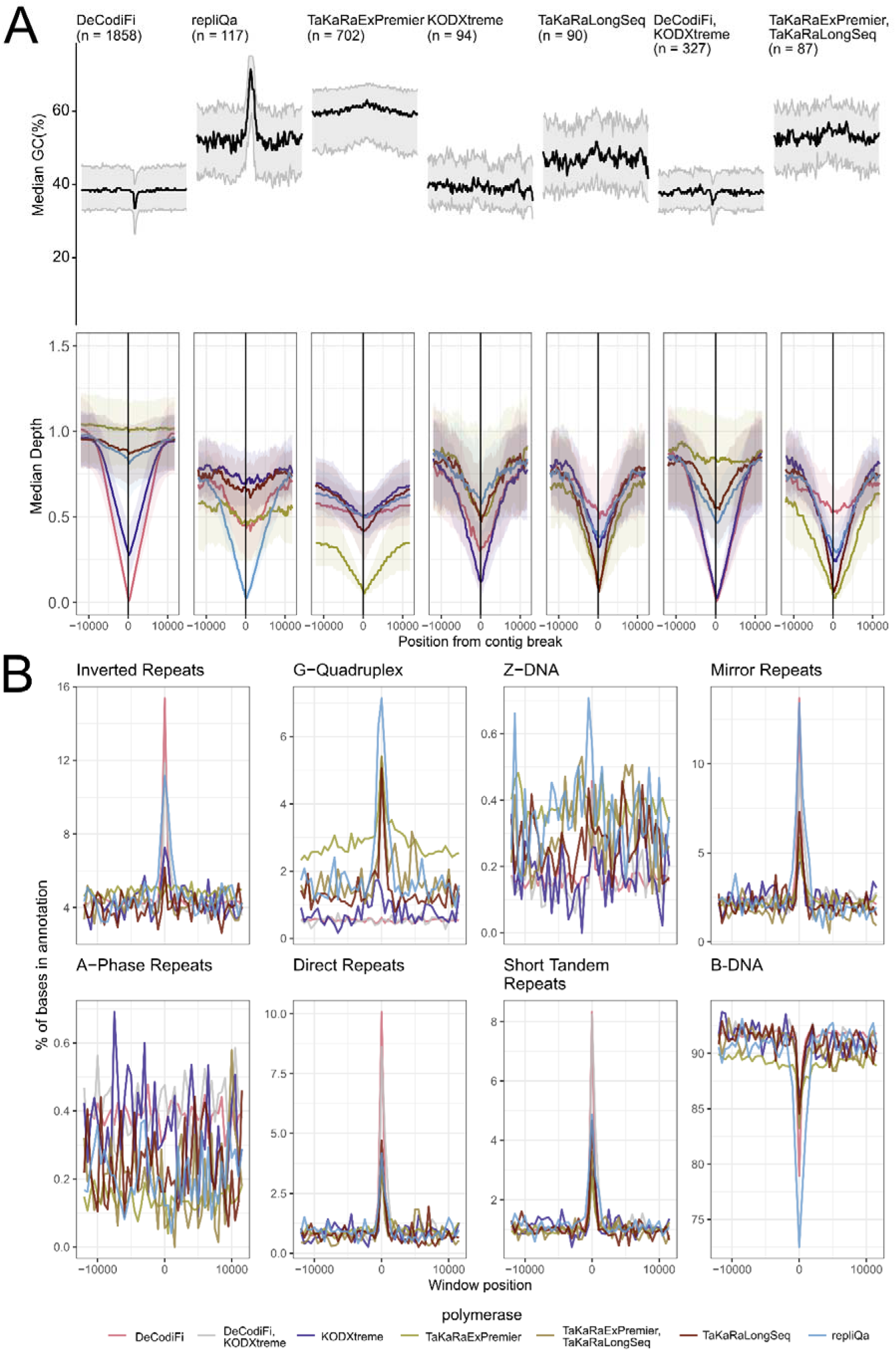
Coverage dropouts and non-B-DNA enrichments in assembly gaps. (A) Median GC-content and normalized coverage centered around all polymerase-specific as well as TaKaRaExPremier-TaKaRaLongSeq and KODXtreme-DeCodiFi shared assembly gaps. Each assembly gap is centered at 0. The GC-content of non-overlapping 200 bp (top) and median per-polymerase normalized coverage (bottom) is shown for 12,000 bp up- and downstream of the aggregated assembly gaps. The number of independent assembly gaps is shown on top of each plot, shaded areas demarcate the 25% and 75% quantile of each window for both read coverage and GC content. (B) Non-B-DNA annotations in assembly gap windows as in (A). For each non-B-DNA category, the percentage of bases assigned to the category in non-overlapping 200 bp windows across aggregated assembly gap windows is shown. The bottom-right plot shows the proportion of B-DNA, defined as DNA not classified as any non-B-DNA structure. Because repeat motifs can be assigned to multiple categories, the cumulative percentage of bases annotated within a given window may exceed 100%.

GC content influences these patterns, with repliQa, TaKaRaExPremier and, to a lesser extent, TaKaRaLongSeq often showing assembly gaps in high-GC regions, whereas DeCodiFi tends to have assembly gaps in low-GC regions (Figure 5A). As illustrated in Figure 4B, we note that complex patterns also exist, in which two or more polymerases exhibit assembly gaps. Overall, this analysis demonstrates that all polymerases fail to amplify certain genomic positions, implying that the increased contiguity and gene completeness of multi-polymerase based assemblies arise from polymerases complementing each other and providing coverage in regions where others drop out.

To get more insights into the DNA sequence motifs that hamper amplification, we annotated non-B-DNA motifs that can form noncanonical DNA structures in the *Carollia* reference assembly with non-B-gfa [43], and intersected these motifs with polymerase-specific assembly gaps (Figure 5B). We found that the regions around DeCodiFi assembly gaps are enriched for different non-B-DNA repeat classes (inverted repeats, mirror repeats, direct repeats, short tandem repeats), whereas assembly gaps only observed for TaKaRaExPremier, repliQa and TaKaRaLongSeq show an enrichment of G-quadruplexes, in line with elevated GC-content around their gaps (Figure 5A). However, whether dropouts are driven by high GC-content or G-quadruplexes remains unclear, as G-quadruplexes are naturally rich in GC. Overall, while the underlying mechanisms require further study, polymerase assembly gaps exhibit notable differences in non-B-DNA motifs.

### Combinations of polymerases also improve human genome assemblies

Based on our previous findings that KODXtreme-based amplification generated more contiguous and complete assemblies for the human HG002 sample than the long-range polymerases used in the original ULI protocol [24], KODXtreme was subsequently adopted in PacBio’s Ampli-Fi library preparation workflow [44]. To assess whether amplification with additional polymerases could further improve human genome assembly quality, we prepared amplification-based libraries with repliQa, TaKaRaLongSeq and DeCodiFi and sequenced one SMRT Cell per polymerase. We also performed an independent KODXtreme amplification, which resulted in highly consistent assembly quality (Figure 3C-D, Figure S10, Table S12).

Assemblies generated from individual polymerase datasets after downsampling to equal coverage showed that neither repliQa nor DeCodiFi outperformed KODXtreme in terms of contiguity (contig N50 1.68 Mb and 110 kb vs. 4.24 Mb for KODXtreme) or gene completeness (96.47% and 79.59% vs. 98.96% for KODXtreme; Figures 3, S10, Table S13), contrasting our observations in *Carollia* and the other test species. TaKaRaLongSeq, however, produced the most contiguous and complete HG002 assembly (contig N50 8.17 Mb, compleasm completeness 99.27%).

Consistent with our results for the other species, combining reads from different polymerase amplifications further improved assembly quality also for the human genome. For example, combining reads from the two KODXtreme amplifications with TaKaRaLongSeq reads increased contig N50 from 8.17 Mb to 12.16 Mb and gene completeness from 99.27% to 99.39% compared with the TaKaRaLongSeq-only assembly (Figure 3C-E, Table S13).

### Variant calling accuracy of amplified reads is comparable to non-amplified reads at matched coverage

The Ampli-Fi library preparation protocol can be applied in clinical settings to characterize genomic variants in human samples that contain only little HMW DNA. We therefore assessed whether improved assembly contiguity and completeness translate into enhanced variant calling performance, as measured by key metrics for clinical utility such as precision, recall, and F1 scores of single nucleotide variants (SNVs), Insertions and Deletions (Indels), Tandem Repeats (TRs), and Structural Variants (SVs). To this end, we retrieved existing HG002 ULI datasets as well as a non-amplified HiFi read dataset as a gold standard (Table S14), and mapped reads downsampled to equal coverage to the hg38 human reference with minimap2 [45]. We inferred SNVs and InDels with DeepVariant [46], Tandem Repeats with trgt [47], and larger SVs with sniffles2 [48]. We then benchmarked our variant calls against the (i) Genome in a Bottle Consortium (GIAB) reference datasets HG002 v4.2.1 for SNV and InDels, (ii) the reference dataset HG002 Tier1 SV v0.6 for SVs, and (iii) the reference dataset HG002_TR_v1.01 for TRs.

TaKaRaLongSeq- and KODXtreme-amplified reads achieved higher SNV, InDel SV and TR F1 scores than reads generated using the original ULI protocol [27], and all amplified reads outperformed those generated with dMDA (Tables S16, S18). TaKaRaLongSeq-amplified reads outperformed reads generated with KODXtreme, repliQa or DeCodiFi. Notably, TaKaRaLongSeq-amplified reads achieved F1 scores close to those of non-amplified Revio-sequenced GIAB reads (SNV 0.999 vs. 0.999; InDel 0.980 vs. 0.990, SV 0.932 vs. 0.944, TR 0.949 vs. 0.964, Table S15).

To investigate whether polymerase-amplified reads can represent variants in challenging genomic regions, we additionally benchmarked SNVs, InDels and SVs against the HG002 Challenging Medically Relevant Gene (CMRG) v1.00 reference dataset [49]. Polymerase-amplified reads reached F1 scores comparable to non- amplified reads, with TaKaRaLongSeq and KODXtreme even slightly outperforming non-amplified reads in SV calling (SV 0.906 vs. 0.904; Figure 6B, Table S15). Of the amplified reads, TaKaRaLongSeq showed the highest F1-scores for SNVs and Indels, whereas KODXtreme yielded the highest F1 score for SVs (Figure 6B). To further assess the robustness of our results, we used an additional reference dataset for small variants and SV (v5.0q [50]), another variant caller longcallD [51] and the benchmarking tool aardvark [52] (see methods). In line with our previous results, amplified reads from TaKaRaLongSeq, KODXtreme and repliQa achieved variant-calling accuracies only slightly lower than non-amplified reads for SNVs, InDels and SVs (Tables S17-18).

**Figure 6:**
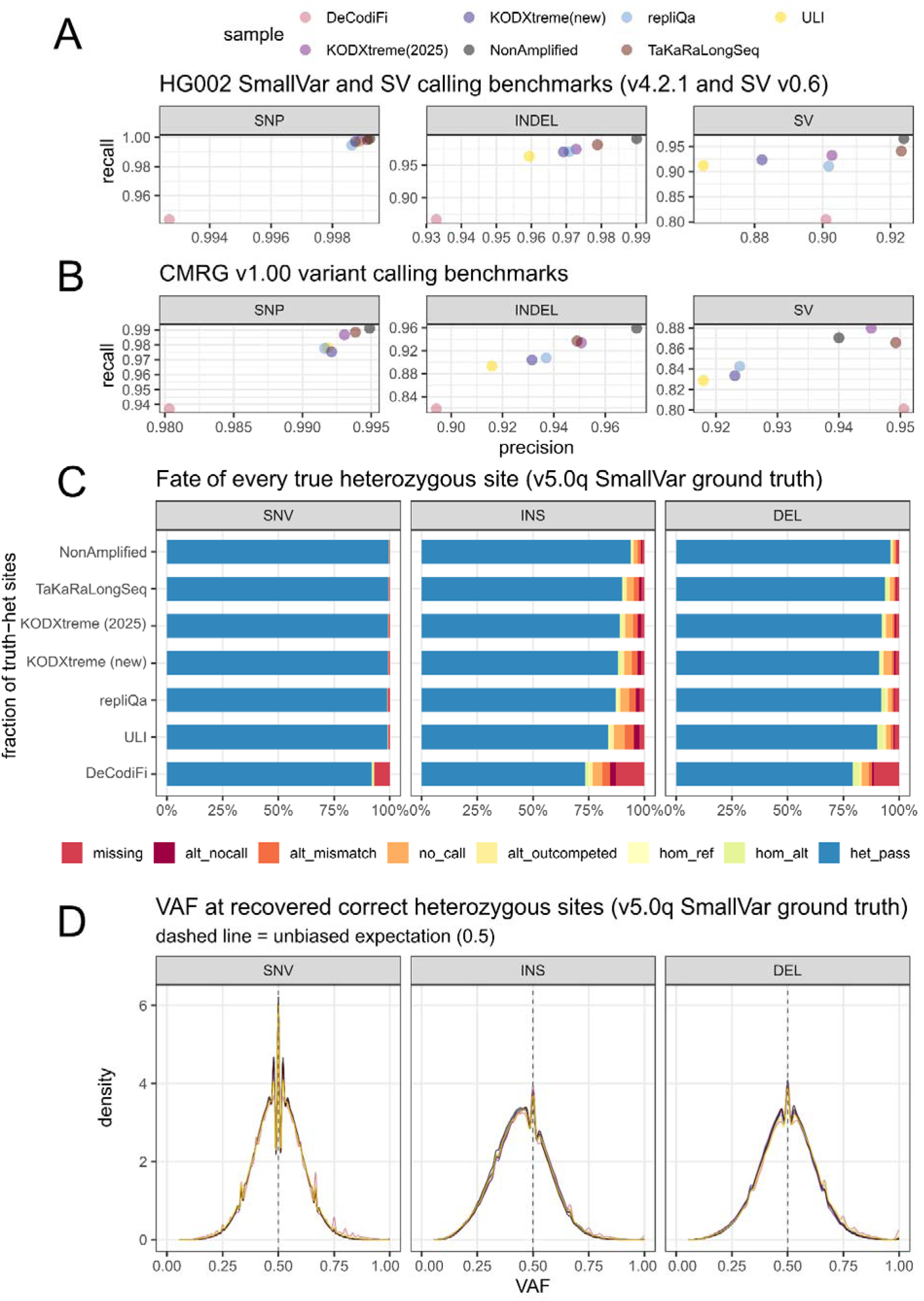
Variant calling benchmarks of polymerase-amplified reads sequenced from the HG002 human sample compared with a non-amplified control. (A) Precision and Recall for ground truth variants (HG002 v4.2.1 and SV v0.6) of six amplified HG002 read datasets and a non-amplified control, downsampled to the coverage of the smallest dataset (KODXtreme(new), ∼25X). (B) Precision and Recall for ground truth variants in challenging medically relevant gene regions (HG002 CMRG v1.00) of six amplified HG002 read datasets and a non-amplified control, downsampled to the coverage of the smallest dataset (KODXtreme(new), ∼25X). (A),(B) SNP = Single nucleotide polymorphism, InDel = small Insertions/Deletions, SV = Structural Variants. (C) Recovery of true heterozygous small variants from the comprehensive ground truth HG002 v5.0q of the same read sets as in (A),(B). Reads are sorted by the number of correctly recovered variants. missing = variant not present in vcf, alt_nocall = at a true position, a different variant was marked NoCall, alt_mismatch = at the position, a different alternative allele was called, no_call = a true alternative variant was marked NoCall, alt_outcompeted = multi-allele call, where a variant other than the true alternative variant was selected by the caller, hom_ref = true alternative allele was marked as RefCall in the vcf, hom_alt = the true alternative allele was called homozygous, het_pass = a true heterozygous variant was recovered and marked “PASS” by the caller. (D) Variant Allele Frequency (VAF) distribution of true heterozygous variants recovered from the same read sets as in (C). To make SNVs, Insertions and Deletions comparable, density distributions are plotted. Line color represents the read dataset. (C),(D) SNV = Single nucleotide variants, INS = Insertions, DEL = Deletion. We note that the frequency of alternative insertion alleles appears to be systematically underestimated for both amplified and non-amplified reads, potentially caused by reference-bias, which affects insertions the most [53].

In human genomics, an unbiased representation of heterozygous variants in sequencing data is paramount for any clinical application. To ascertain allelic imbalance or dropout in amplified read datasets, we determined how many true heterozygous SNVs and InDels (ground truth: v5.0q smallvar) were correctly genotyped for each downsampled benchmarked sample, and whether the fraction of reads supporting the alternative allele significantly deviates from the expectation of 0.5. Mirroring our variant calling benchmarks, TaKaRaLongSeq, KODXtreme and repliQa recovered more true heterozygous variants than the best ULI dataset, approaching the performance of non-amplified reads (Figure 6C, Table S19). Likewise, variant allele frequency distributions of SNVs and InDels closely resemble those of non-amplified reads (Figure 6D) and only a small fraction of correctly recovered heterozygous alleles showed VAFs significantly different from 0.5 (maximum: DeCodiFi: 949 [0.036%] true sites deviating from 0.5; Table S19), indicating that amplification does not introduce a skewed allele representation.

We further asked whether amplified reads are amenable to variant phasing. To this end, we used HiPhase [54] to phase variants into haplotype blocks and counted the number of confidently phased variants. We note that read length is likely the dominating factor for phasing accuracy, and polymerase-amplified reads are on average considerably shorter than non-amplified reads (∼10 kb vs. ∼16 kb). Still, at matched coverage, a similarly high fraction of variants could be phased in amplified and non-amplified read sets (e.g. TaKaRaLongSeq vs. non-amplified: 98.0% vs 98.2%, Table S20), consistent with our variant calling benchmarks and the allelic imbalance analysis.

Together, these results indicate that optimized amplification-based HiFi sequencing protocols can enhance variant detection accuracy in low-input human samples compared with the Ampli-Fi protocol, while introducing minimal allelic imbalance or allele drop-out. Our findings suggest that these protocols may be applicable to challenging clinical materials such as dried blood spots, cancer biopsies, or saliva samples [27,28], thereby increasing their potential utility in medical research and diagnostics.

### Improved reference genomes of Elysia and Carollia

Finally, in addition to the new *Tetrodontophora* genome described above, we pooled new HiFi reads generated in this study to improve the reference genomes of *Elysia* and *Carollia*. To scaffold the assemblies, we generated Arima Hi-C data for Carollia (∼90X coverage) and used existing Hi-C data for *Elysia* [23]. For *Elysia*, we generated and curated a new primary scaffolded genome, achieving improved contig N50 (2.31 vs. 1.92 Mb) and scaffold N50 (45.5 vs. 41.8 Mb, Figure S11) values compared with the existing reference genome [23]. For *Carollia*, we generated a curated, haplotype-resolved, chromosome-level reference genome that improves contig N50 (haplotype 1 49.82 vs. 31.70 Mb) and scaffold N50 (haplotype 1 278.72 vs. 240.44 Mb, Figure S12) compared with the existing PacBio CLR-based reference [39].

## Discussion

Our benchmark comparing eleven long-range PCR polymerases across organisms representing three evolutionarily divergent eukaryotic lineages demonstrates that genome assembly quality varies substantially depending on the polymerase used for amplification. With the exception of LongAmpTaq and TaKaRaGXL, which showed low performance for *Elysia* and *Carollia* respectively, polymerase performance was generally consistent across the three species. This concordance allowed us to identify trends that may have general applicability beyond individual taxa.

We identified five polymerases that consistently outperformed the other enzymes across the three test species. Among these, single-polymerase assemblies generated with repliQa often produced the highest-quality assemblies, consistent with a recent benchmark in yeast [31]. Notably, for *Tetrodontophora* and *Carollia,* assemblies generated with repliQa and TaKaRaLongSeq were markedly better than those generated with KODXtreme, the polymerase currently employed in PacBio’s Ampli-Fi protocol. TaKaRaExPremier also outperformed KODXtreme for *Tetrodontophora*. Together, this indicates that the current Ampli-Fi protocol can be improved.

Across all eleven polymerases, the primary reason for differences in assembly quality is their ability to amplify GC-poor and GC-rich DNA. Whereas the top-performing polymerases exhibited a relatively low bias, other enzymes showed severe difficulties in amplifying regions with <25% or >70% GC content. Although such extreme GC regions often make up only a small fraction of the genome, systematic coverage reduction or dropout in these regions had a disproportionate negative impact on assembly contiguity and completeness. Polymerase-specific dropout regions were also enriched in regions predicted to form noncanonical DNA structures. Such regions have been associated with systematically poor long-read sequencing performance of bird dot chromosomes [55] and *Drosophila* Y-linked genes [56]. This is consistent with altered polymerization kinetics and sequencing performance in such regions across PacBio, Illumina, and Oxford Nanopore platforms [57,58], and suggests that noncanonical DNA structures are an additional, polymerase-dependent source of amplification bias.

Although none of the polymerases included in our benchmark was free of amplification bias, we observed that combining two or more polymerases often yielded superior assemblies. By analyzing read coverage around polymerase-specific assembly gaps, we demonstrate that this improvement arises from polymerase complementarity, where dropouts of one polymerase are compensated by sequencing reads generated by others. While stochastic effects during DNA extraction, amplification or library preparation, and limited library complexity may also contribute to performance variability, these effects were minor compared to the impact of using multiple polymerases. Taken together, our results indicate that assembly quality can be most effectively improved by performing multiple amplifications using different polymerases.

Our evaluation of variant-calling performance showed that reads amplified with the best-performing polymerases achieved accuracy comparable to non-amplified reads, with minimal allelic imbalance and reliable representation of challenging, medically relevant regions of the human hg38 genome. In particular, our results identify TaKaRaLongSeq as a promising candidate to replace KODXtreme in an improved, single-polymerase Ampli-Fi protocol for human whole-genome sequencing.

Our general recommendation for generating a long-read-based genome assembly of a new species is to use multiple polymerases, ideally selecting three enzymes that show complementary performance (TaKaRaLongSeq or TaKaRaExPremier, repliQa, and KODXtreme or DeCodiFi; Table S21). By using amplification primers with distinct barcodes, the resulting libraries can be pooled in equal proportions and sequenced together, while retaining the ability to separate the reads by polymerase. This allows evaluating assembly performance at matched sequencing coverage and facilitates the selection of the best-performing polymerase combination.

Our study provides a general framework for benchmarking newly developed polymerases and extending such analyses to additional taxonomic groups. This will be essential to further assess the generality of the proposed strategy across a larger spectrum of eukaryotic genome diversity and to identify optimal polymerase combinations for particular groups. To facilitate the computational aspects of future benchmarks, we provide the polymeval pipeline (https://github.com/casparbein/polymeval) [59], which automates contig assembly with or without downsampling from individual or combined polymerase datasets, as well as variant calling benchmarking for HG002 samples, and generates standardized assembly quality metrics and visualizations.

## Conclusion

In summary, our results demonstrate that genomic DNA amplification enables long-read sequencing of samples such as ethanol-preserved specimens, specimens containing potential polymerase-inhibiting metabolites, minimally invasive biopsies, and small-bodied organisms, which do not meet the gold-standard requirement of sufficient amounts of flash-frozen tissue. Using optimized amplification strategies, we routinely generated assemblies exceeding 1 Mb contig N50 and 90% BUSCO completeness, representing a substantial improvement over short-read–based approaches. Although further validation across additional taxa is required, our data show that the Ampli-Fi protocol polymerase KODXtreme can be outperformed by other enzymes for certain species, and that the combined use of multiple polymerases (as in the original ULI protocol) regularly yields superior assemblies. We anticipate that adoption and customization of amplification-based long-read protocols, including substituting or combining polymerases within the Ampli-Fi framework, will provide the community with a powerful toolkit for generating high-quality reference genomes from challenging samples.

## Competing interests

J. A. L. and I. M. are employees and shareholders of Pacific Biosciences. Other authors have no competing interests.

## Acknowledgments

This work was supported by the Leibniz Association’s Competition Procedure (K419/2021), the LOEWE-Centre for Translational Biodiversity Genomics (TBG) funded by the Hessen State Ministry of Higher Education, Research and the Arts (LOEWE/1/10/519/03/03.001(0014)/52), and the European Research Council (ERC) under the European Union’s Horizon 2020 research and innovation programme (grant agreement No. 101118919). We thank Gregory Young, Aaron Wenger and Jonas Korlach from PacBio for the human sample runs of DeCodi-Fi, KODXtreme, TaKaRaLongSeq and repliQa. We furthermore thank Christoph Sinai for assistance in running the analysis on a HPC, Andreas Kaut for helping with sample preparation and Bruno Hüttel for his help in setting up the qPCR experiments. We also thank the Bioscientia Institut für Medizinische Diagnostik GmbH (Ingelheim, Germany) for providing the PacBio SMRT sequencing service on the PacBio Revio platform.

## Data and Code Availability

Polymerase-stratified, amplified reads for *Tetrodontophora*, *Carollia* and *Elysia* are available on NCBI under Bioproject (PRJNA1544338). The haplotype-resolved *Tetrodontophora* and *Carollia* and scaffolded *Elysia* assemblies are likewise available on NCBI under the same Bioproject ID. Amplified HG002 reads are available on the PacBio cloud (https://downloads.pacbcloud.com/public/2026Q2/Ampli-Fi/). All scripts used in the analysis are available as a pipeline on github at https://github.com/casparbein/polymeval [59], distributed under the MIT-License.

## Materials and Methods

### Polymerase selection

Long-range PCR amplification of long (>1 kb) sequences for LRS and specifically for HiFi sequencing is not yet a widely adopted approach. There are theoretically dozens of polymerases that could potentially be tested. After careful consideration and literature research, we opted to test polymerases which are designed to amplify long fragments according to the manufacturers’ protocol (Table 1). The polymerases we chose generally outperformed other polymerases in previous experiments, mostly for amplicon or short-read sequencing (SRS, Table S1). We and others have previously employed seven of the eleven polymerases for PacBio HiFi LRS (Table S1) [24,31]. Nevertheless, to our knowledge, this is the first study that tests these polymerases on a range of eukaryotic genomes with estimated sizes between ∼700 Mb and ∼2.5 Gb.

### Sample sources

Specimens of *Tetrodontophora bielanensis* were collected close to Ostritz-Marienthal, Saxony, Germany in April and June of 2025 (coordinates: 50.99041°N, 14.90080°E). Specimens of *Elysia timida* were collected in Cadaqués, Girona, Spain, in May 2023 (coordinates: 42.285173, 3.296461). Access to genetic resources of *Elysia*, including associated benefit-sharing arrangements and compliance with applicable regulations, was granted under the permit numbers ESNC 205 and ESNC 285. Skin and liver tissue were collected from one male individual of *Carollia perspicillata* from a captive breeding colony in Frankfurt, Germany. Tissues were collected immediately after euthanization and directly stored in pure ethanol. After sampling, these samples were transported on dry ice and moved to −80°C for long-term storage. The human DNA sample was obtained from the NIGMS Human Genetic Cell Repository at the Coriell Institute for Medical Research: HM24385 (NIST ID: HG002).

### DNA extraction

High-molecular-weight (HMW) genomic DNA (gDNA) of *Elysia* was extracted from one whole individual according to the protocol of [60]. For *Carollia,* HMW gDNA was extracted from approximately 19 mg of pestle-homogenized liver tissue following the protocol “Extracting DNA from animal tissue using the Nanobind® PanDNA kit - Procedure & checklist” (102-574-600 REV04 AUG2024) including a preceding ethanol removal step. This step was done according to the “Nanobind® PanDNA kit - Guide & overview” (103-394-800 REV04 OCT2024), but with reduced volumes. Additionally, HMW gDNA was extracted from skin tissue (3 mm biopsy) of the same *Carollia* individual according to the protocol of [60] with a preceding ethanol removal step as for the liver tissue. HMW gDNA of *Tetrodontophora* was prepared from one fresh-frozen animal using a modified version of the QIAGEN MagAttract HMW DNA Extraction Kit protocol (see the Supplementary Material for more details).

The DNA concentration of *Elysia* and *Carollia* was measured using the QuantiFluor® ONE dsDNA System on the Quantus™ Fluorometer (Promega Corporation). The Qubit Fluorometer with the Qubit 1X dsDNA HS Assay-Kit (Thermo Fisher Scientific) was used to quantify the DNA of *Tetrodontophora*. For all samples, DNA fragment length was assessed using the Femto Pulse System with the Genomic DNA 165 kb Kit (Agilent Technologies).

### PacBio Ampli-Fi SMRTbell library preparation

For *Elysia*, *Carollia* (liver tissue) and *Tetrodontophora*, PacBio Ampli-Fi SMRTbell libraries were prepared using a modified version of the Ampli-Fi protocol (for more details see Supplementary Material). For each of the three species, the long-range PCR step during library preparation was performed twice with each of the initial ten polymerases (Tables S1, S22, S23), each time using an aliquot of the same sheared and amplification adapter-ligated DNA. The same DNA was used beforehand to determine the optimal number of PCR cycles for each polymerase-species combination via qPCR to avoid overamplification (Table S24). Following long-range PCR, barcoded polymerase-specific PacBio Ampli-Fi SMRTbell libraries were prepared using the amplified DNA. To minimise bias, all PacBio Ampli-Fi SMRTbell library preparations for each species were performed in parallel. Regarding *Tetrodontophora*, the libraries were equimolarly pooled in groups of ten (two pools in total) directly after barcoding and the final library pools were size selected with AMPure PB Beads to remove < 5 kbp fragments. For *Elysia* and *Carollia* (liver tissue), the individual final barcoded libraries were size selected with AMPure PB Beads to remove < 3 kbp fragments and then pooled equimolarly in groups of five (four pools per species, eight in total). To include an eleventh polymerase, TaKaRaLongSeq, in the initial benchmark for *Elysia* and *Tetrodontophora*, two additional PacBio Ampli-Fi SMRTbell libraries were prepared per species (PCR replicates) from the same sheared DNA as before. After library preparation, the four final libraries (two per species, each with a species-specific barcode) were equimolarly pooled and sequenced on a single SMRT Cell.

Five additional PacBio Ampli-Fi SMRTbell libraries were individually prepared with DNA from skin tissue of the same *Carollia* specimen (for more details see Supplementary Material). This was done using the modified Ampli-Fi protocol individually combined with each of the following polymerases: TaKaRaLongSeq, TaKaRaExPremier, repliQa, KODXtreme, and DeCodiFi. Regarding the polymerases repliQa and DeCodiFi, long-range PCR (including the amplification adapter ligation step beforehand) had to be performed twice, after which the amplified DNA was pooled to provide an adequate quantity of input DNA for the subsequent SMRTbell library preparation step. The final libraries were size selected using a 3 kbp cut-off with AMPure PB Beads for TaKaRaLongSeq and a 5 kbp cut-off with the BluePippin for the other four polymerases. For the human sample SMRTbell library preparation for HiFi sequencing was performed according to the PacBio Ampli-Fi protocol individually combined with each of the following four polymerases: repliQa, TaKaRaLongSeq, KODXtreme and DeCodiFi (for more details see the Supplementary Material).

### HiC - Chromatin conformation capture

Chromatin conformation capture and library preparation for *Tetrodontophora* and *Carollia* was performed using the “Arima High Coverage HiC Kit - User Guide for Animal Tissue” (A160162 v01, November 2021) and the “Arima Library Prep Module - User Guide: Library Preparation for the Arima HiC+ for Genome Wide HiC Kit” (A160432 v02, May 2023), respectively. For *Tetrodontophora*, the low-input protocol was applied to a new individual from the same collection site, followed by an 11-cycle PCR during library preparation. For *Carollia*, the standard-input protocol was used on approximately 28 mg of ethanol-preserved kidney tissue from the same individual, followed by a 10-cycle PCR during library preparation. For both species, the DNA concentration and DNA fragment length were assessed using the Qubit Fluorometer with the Qubit 1X dsDNA HS Assay-Kit (Thermo Fisher Scientific) and the TapeStation 4150 with the D1000 and D5000 ScreenTape (Agilent Technologies), respectively.

### Sequencing

For the initial benchmark with the polymerases involving *Elysia*, *Carollia* (liver tissue), and *Tetrodontophora*, a total of 11 PacBio Ampli-Fi SMRTbell library pools were sequenced. For the subsequent second experiment focusing on the “best” five polymerases using *Carollia* (skin tissue), each of the five polymerase-specific libraries was run on a single Revio SMRT Cell. The same was done for the four polymerase-specific libraries for the human sample. Annealing of sequencing primers, binding of sequencing polymerase, and purification of polymerase-bound SMRTbell complexes were performed using the Revio polymerase kit (PacBio) or SPRQ chemistry for the human samples. The libraries were loaded at an on-plate concentration of 200-300 pM each by adaptive loading. A total of 20 Revio SMRT Cells were sequenced. The Arima HiC libraries were sequenced on the NovaSeq X Plus platform at Novogene (UK) using a 150 bp paired-end sequencing strategy, with an expected output of 23 Gb (*Tetrodontophora*) and 90 Gb (*Carollia*) and an insert size of 350 bp.

### Flow Cytometry

The genome size of *Tetrodontophora* was estimated following a flow cytometry (FCM) protocol with propidium iodide-stained nuclei described by [61]. Two individuals (one male, one female) were each chopped with a razor blade in a Petri dish. Ice-cold Galbraith buffer (2 mL) was used as the suspension medium. The internal reference standard was a separate suspension of Chicken Erythrocyte Nuclei (Biosure, 1C = 1.2 Gb). The three suspensions were filtered through a 42-μm nylon mesh, stained with the intercalating fluorochrome propidium iodide (PI, Thermo Fisher Scientific, final concentration 25 µg/mL), and treated with RNase A (Sigma-Aldrich, final concentration 250 µg/mL). The mean red PI fluorescence signal of stained nuclei was quantified using a Beckman-Coulter CytoFLEX flow cytometer with a solid-state laser emitting at 488 nm. Fluorescence intensities of at least 10,000 events per measurement were recorded. We used the software CytExpert 2.3 for histogram analyses. After measuring the suspensions of *Tetrodontophora* and the internal reference standard separately, the samples were each mixed with the standard. The total quantity of DNA per nuclei of each *Tetrodontophora* individual was calculated as the ratio of the mean red fluorescence signal of the 2C peak of the stained nuclei of *Tetrodontophora* divided by the mean fluorescence signal of the 2C peak of the stained nuclei of the reference standard times the 1C amount of DNA in the reference standard. The two *Tetrodontophora* individuals were each measured as triplicate, with the three measurements carried out on three different days to minimize possible random instrumental errors. The average of these three measurements was then calculated to estimate the genome size (1C) of each *Tetrodontophora* individual. The value of the robust coefficient of variance (rCV) was approximately 5% or less indicating very little spread in fluorescence intensity between runs or samples for a given cell population.

### Read processing

For each polymerase amplification of the first experiment, we obtained reads from two different pools. After preliminary analysis, we deemed single-pool read output too low to draw meaningful conclusions and therefore merged same-polymerase reads per species prior to further processing. We then removed adaptors and demultiplexed sequencing reads of *Carollia*, *Elysia* and *Tetrodontophora* per polymerase read set with lima v2.13.0 (https://github.com/pacificbiosciences/barcoding/). For all reads analysed in this study, we removed PCR duplicates with pbmarkdup v.1.2.0 (https://github.com/PacificBiosciences/pbmarkdup). For each same-polymerase read set, we then computed basic read summary statistics such as number of reads, number of sequenced bases, read N50 and average read length with seqkit v2.14.0 [62] and plotted read length histograms with a custom R script [63].

### In-silico assembly experiments

To ensure a reproducible experimental design for comparing polymerase-specific assemblies across organisms, we developed ‘polymeval’, a pipeline to evaluate polymerase amplification and sequencing performance for assembly. Polymeval is implemented in snakemake v9.27.1 [64] and available on github (https://github.com/casparbein/polymeval)[59].

The pipeline consists of several modes: In standard mode, it computes primary polymerase-library specific assemblies for each organism with all available data per library with hifiasm v0.25.0 [34] and strict haplotig purging (parameter –l 3) to compare assembly contiguity as measured by contig N50 and assembly span. Since different available assembly tools have been shown to produce relatively more contiguous and complete assemblies depending on sequencing technology, input read characteristics and organism [65], we initially also computed assemblies with flye v2.9.6 [66] and LJA v0.2 [67], but LJA failed to produce assemblies for more than a third of the reads we passed, and the flye assemblies were always less contiguous and complete than the hifiasm assemblies (Table S25), prompting us to continue all following assemblies with hifiasm. For the first *Carollia* amplification experiment, we added the expected haploid genome size (---hg-size 2g) because we saw that several assemblies were otherwise insufficiently purged. The pipeline also puts out corrected reads from hifiasm’s internal error correction module (--write-ec). The proportion of corrected bases per sequenced base is then calculated with hifieval v0.4.0 [38]. Subsequently, polymeval runs compleasm v0.2.9 [35] with the mammalia_odb12 (for *Carollia* and human), arthropoda_odb12 (for *Tetrodontophora*) and mollusca_odb12 (for *Elysia*) orthologue sets to measure assembly completeness and detect potential unpurged haplotigs, which would be visible by an excess of duplicated orthologues. All results are then plotted by custom R scripts [63]. Since sequencing output as measured in number of sequenced bases differed extensively between polymerase-libraries, input reads have to be downsampled for a fair comparison. Therefore, polymeval has a “downsample” mode where it downsamples all libraries to the number of sequenced bases found in the library with the smallest output (sequenced bases) with Rasusa v5.1.0 [68]. For *Elysia*, LongAmpTaq provided less than one third of the output of the next smallest library and was therefore discarded for these experiments. Next, polymeval assembles the downsampled libraries and assesses assembly completeness as described above.

Furthermore, we designed several downsampling-and-combine assembly experiments to investigate whether repeated amplifications with the same library or a combination of different libraries lead to more contiguous assemblies, using polymeval’s ‘combine’ mode. First, for each species we defined a target coverage *t* as the 0.5X of the smallest read dataset of that species (excluding LongAmpTaq in *Elysia*) and downsampled all read datasets to that coverage. We then built all possible pairwise combinations, leading to 36 pairs for *Carollia* (skin tissue), 45 pairs in *Elysia* and *Carollia* (liver tissue) and 55 pairs in *Tetrodontophora*, and looked at assembly contiguity and completeness (Tables S6, S11).

Next, based on the results from downsampling and combination experiments as well as GC dropouts and coverage heterogeneity, we chose five polymerases to investigate further (repliQa, TaKaRaExPremier, TaKaRaLongSeq, KODXtreme, DeCodiFi). We asked whether combinations of more than two polymerases as tested above would increase assembly quality even more. Thus, for each species and sequencing experiment (two for *Carollia*), we again defined *t* as the smallest output among those five polymerases (for the first *Carollia* experiment, we used TaKaRaGXL instead of TaKaRaLongSeq), and then downsampled all read datasets to 1*t*, 0.5*t*, 0.33*t*, 0.25*t* and 0.2*t*. We then built all combinations such that overall input for hifiasm was 1*t,* leading to 31 assemblies per species and experiment (Figure 3, Tables S8-S9). For the HG002 sample, we used the same approach with TaKaRaLongSeq, KODXtreme, DeCodiFi and repliQa, additionally including a previously sequenced KODXtreme dataset [24] (Table S13).

Since batch effects might influence coverage heterogeneity by random over- or undersampling of genomic regions in a given amplification, we next asked whether combinations of different polymerases, and/or combinations of the same polymerase from different amplification runs improve assembly quality. Because we had two independent amplification experiments for *Carollia* for four polymerases, the subsequent experiments were performed with *Carollia* reads amplified with DeCodiFi, KODXtreme, TaKaRaExPremier and repliQa. For each polymerase, we defined a target coverage *t*, set to 2X the input nucleotides of the first experiment dataset. We then downsampled reads from the second sequencing experiment to 1*t* and 0.5*t* (Table S10), assembling all combinations with *t* input. For example, we sequenced 28 Gb KODXtreme reads in the first experiment, and *t* is 2X 28 Gb or 56 Gb. Therefore, we downsampled KODXtreme reads from the second experiment to 0.5*t* (28 Gb) and 1*t* (56 Gb). We then created two assemblies: 0.5*t* KODXtreme (first experiment, 28 Gb) + 0.5*t* KODXtreme (second experiment, 28 Gb), and 1*t* KODXtreme (second experiment, 56 Gb). Finally, given four polymerase, we obtained eight assemblies.

When performing in-silico downsampling and combining reads, we note that the specific ranking of assemblies based on contiguity and completeness can change slightly depending on the chosen pseudorandom seed. Importantly, the polymerases we identified as generally more useful for long-read sequencing outperform the other polymerases independent of the downsampling seed. Likewise, where assemblies computed from combined read sets ranked higher than single-polymerase assemblies, they did so consistently (5 seeds were tested, not shown).

### *Tetrodontophora* and *Carollia* haplotype-resolved assembly

We assembled the genome of *Tetrodontophora* and *Carollia* using a semi-automated assembly pipeline that follows the best practices proposed by the Earth BioGenome Project. In brief, we trimmed PacBio HiFi-CCS long reads and Hi-C Illumina short reads using Cutadapt v5.0 [69] and Fastp v0.23.4 [70], respectively, to guarantee high-quality input data for assembly. FastQC v0.12.1 (https://github.com/s-andrews/FastQC) was used for quality control of the HiC reads, and Nanoplot v1.44 [71] was used to evaluate the long-read characteristics. K-mer-based profiling was conducted using FastK v1.1.0 (https://github.com/thegenemyers/FASTK). We estimated genome size, heterozygosity, and repetitive content with GenomeScope2 v2.0.1 [72], used SmudgePlot v0.4.0 [72] to infer ploidy levels through the K-mer distribution analysis, and used KatGC v1.1.2 [73] to investigate possible GC content deviations in relation to K-mer distribution.

A phased genome assembly was generated with Hifiasm v0.25.0 [34] using both PacBio HiFi and Hi-C reads as input. BandageNG v2025.6.1 [74] was used to investigate the assembly graphs. The fasta file for each haplotype was generated with GFAstats v1.3.11 [75]. The mitochondrial genome was assembled with MitoHifi v3.2.3 [76]. Putative contaminants were identified and removed using FCS Adaptor v0.5.0 and FCS-GX v0.5.0 [77]. To scaffold the contigs, we mapped Hi-C reads to the phased genome assemblies. Genome indexing and mapping for both haplotypes from decontaminated assemblies were performed using Chromap v0.3.0-r509 [78]. Subsequently, forward and reverse Hi-C reads were aligned separately to each haplotype and sorted with SAMtools v1.21 [79], using default parameters for HiC mapping. Mapping statistics were assessed using SAMtools flagstats. We used PretextMap v0.1.9 (https://github.com/sanger-tol/PretextMap) and PretextSnapshot v0.0.4 (https://github.com/sanger-tol/PretextSnapshot) to build and obtain visual representations of Hi-C contact maps. The automatic scaffolding step, performed using YAHS v1.2.2 [80], linked contigs into longer scaffolds to improve the assembly contiguity. Following automatic scaffolding, we manually curated the genome by remapping the Hi-C reads to the concatenated haplotypes using Chromap, but allowing for the presence of duplicate mappings. The manual curation of each haplotype was conducted in PretextView 1.0.3 (https://github.com/sangertol/PretextView). After curation, we built the haplotype curated sequences using agp-tpf-utils v1.2.3 (https://github.com/sanger-tol/agp-tpf-utils).

### *Elysia* assembly

Primary contigs were assembled with Hifiasm 0.19.8-r603 from HiFi reads only. Subsequently, the primary contigs were filtered for artificial sequences and contamination with FCS-adaptor and FCS-GX 0.5.5 [77], respectively. To do so, fcs.py screen genome was run with the NCBI taxid 154625 and a FCS-GX database built on 2026-07-01. Afterwards, fcs.py clean genome was executed with the primary contigs from hifiasm and the action report from “screen genome” as well as the option “--min-seq-len 1”. The contigs remaining after contamination screening were polished with a workflow containing DeepVariant [46]. The HiFi reads used for assembly were mapped against the contamination filtered contigs with minimap 2.26-r1175 [45] with the preset “-x map-hifi”. Duplicates were removed from the bam file with Picard 3.4.0 (https://github.com/broadinstitute/picard) MarkDuplicates and the option “--REMOVE_DUPLICATES true”. Variants were called with DeepVariant 1.10.0 [46] and the option “--model_type=PACBIO”. Homozygous variants were obtained with bcftools 1.10.2 [81] view and the options -f ‘PASS’ -i ‘GT=“1/1”’. The remaining variants were incorporated into the assembly by bcftools consensus (25171 in total). Scaffolding was done by mapping the Hi-C reads to the polished contigs with chromap 0.3.2 [78] by creating an index (-i) and mapping itself with the options “--preset hic --remove-pcr-duplicates --SAM”. The resulting sam file was sorted by read name (-n), converted to bam format (-O BAM) and indexed with samtools 1.20 [81] sort and faidx respectively.

Scaffolding was carried out with yahs 1.1a-r3 [80] in combination with the polished contigs and the Hi-C mapping bam file. The contact map was stored in hic format with yahs’ tool “juicer pre” and JuicerTools 1.22.01. Visualization and manual curation was carried out in Juicebox 2.16 [82].

The final assemblies were validated for completeness and accuracy. We assessed gene-space completeness using Compleasm v0.2.9 [35] against the arthropoda_odb12, laurasiatheria_odb12 and mollusca_odb12.2 BUSCO database, respectively. Assembly consistency was further evaluated by comparing read-level k-mer distributions to the final assemblies using Merqury.FK (https://github.com/thegenemyers/MERQURY.FK). Finally, assembly metrics and quality parameters were summarized in a snail plot generated by BlobToolkit v4.4.6 [83] and blobtk v0.6.5 [84].

### Assessing polymerase sequencing performance

Summary statistics like the contig N50 value or compleasm completeness can quickly point out systematic issues with an assembly, but the underlying causes, such as read coverage dropouts or chimeric reads (defined as a read with at least two parts originating from different genomic regions) remain hidden. We therefore wanted to further assess potential reasons for divergent assembly quality values. For all analysis, we mapped deduplicated polymerase reads to the respective reference (*Tetrodontophora*: our haplotype 1, *Elysia*: GCA_043644045.1 [23], *Carollia*: [39] with minimap2 v2.30 [45].

To investigate potential chimeric reads, we counted the proportion of primary, non-primary (likely repetitive sequences) and supplementary (chimeric) alignments in each unfiltered bam file and plotted length of primary and supplementary alignments against read length (Figure S7). As we previously discovered, even a few systematic read dropouts can drastically decrease contiguity and completeness [24]. Consequently, we wanted to investigate coverage unevenness, GC-bias (systematic coverage dropouts in genomic regions with high or low GC-content) as well as DNA sequence motifs correlating with read dropout. Thus, we computed sequencing coverage from bam files in 200 bp non-overlapping windows with PanDepth v2.26 [85], filtering out unmapped reads, non-primary alignments and alignments flagged as duplicates. To make coverage values comparable across different sequencing depths, we normalized coverage by dividing per-window coverage by the overall mean coverage. Because we saw that normalized coverage was disproportionally affected by outlier genomic windows with sometimes 10,000 X normalized coverage in *Carollia* mappings (putatively collapsed repeats in the CLR reference assembly), we removed windows with >50 X normalized coverage from the calculations (removing 0.001% of windows). If not otherwise indicated, we ran the analysis as implemented in polymevals ‘reference’ mode.

Since polymerases can differ in the extent in which certain fragments are amplified compared to all other fragments [86], we estimated coverage unevenness following different approaches. Particularly, we computed the genome breadth, which is the percentage of reference bases covered by at least one read [87], the low-coverage index (LCI, [31,88]), defined as the percentage of reference sequence covered by less than or equal 0.5 normalized coverage, and coverage heterogeneity, which we define as the median of all Interquartile Ranges of normalized coverage, computed in 100 kb non-overlapping windows along the reference (following [89]). For even coverage, we expect genome breadth to approach 1 and the LCI and coverage heterogeneity to approach 0. We also plotted the distribution of normalized coverage of 200 bp non-overlapping windows across each polymerase sample, where a single, high mode around 1 indicates a more even coverage and a spread out distribution suggests more uneven coverage.

To investigate the GC-bias of each polymerase, we calculated the average GC-content of 200 bp non-overlapping windows as before. Subsequently, we plotted normalized coverage per polymerase readset against GC content bins by steps of 0.01 and visually inspected the coverage distribution across all GC content bins (Figure 2, Figure S8C). GC-dependent dropouts are apparent if the median normalized coverage in a given GC-content bin approaches 0.

### Characterization of polymerase-specific dropouts on the nucleotide level

To gain more insights into which specific regions affect amplification performance negatively, we investigated polymerase read coverage at contig breaks, the regions up- and downstream of assembly gaps in the respective assemblies. We focused on *Carollia* using the deeper coverage sequencing data generated from skin tissue (polymerases: DeCodiFi, KODXtreme, repliQa, TaKaRaExPremier, TaKaRaLongSeq). Using the polymeval ‘reference’ mode, we mapped the five polymerase-specific assemblies retrieved from a polymeval ‘standard’ run to the *Carollia* reference genome with minimap2 v2.30 [45] in ‘paf’ mode. We then processed each pairwise alignment with the readPaf function from the SVbyEye R package [90], filtering out alignments shorter than 1000 bps and with a mapping quality below 50, and identified contig breaks, as defined by the last position of a query contig aligning to the reference. We added 2kb windows up- and downstream of the breakpoints and joined overlapping breakpoint windows. Next, we filtered breakpoint windows exceeding a size of 20 kb, to ensure that the break is not caused by an overarching misassembly in the reference or query. We further counted the number of breakpoint windows per polymerase assembly and joined breakpoints between two or more polymerase-assemblies. We then added another 10kb up-and downstream of these windows and computed polymerase-specific read coverage, normalized coverage and GC-content as before with PanDepth v2.26 [85]. To visually inspect coverage at contig breakpoints, we plotted contig breaks along the longest reference scaffold and chose 30 regions containing at least two polymerase specific contig breaks, displaying coverage in breakpoint windows (total window size = 24 kb, Figure 4). To generalize dropout patterns in single windows, we aggregated all polymerase-specific breakpoint windows, centering the breakpoint at 0 and displaying median normalized coverage and median GC content of each 200 bp non-overlapping window in the aggregate range, with coverage stratified by polymerase.

Since we discovered polymerase-exclusive characteristics in aggregate breakpoint windows, we wanted to evaluate whether non-canonical three-dimensional DNA folding motifs (non-B-DNA) are enriched at breakpoints and whether different polymerase breakpoints show unequal enrichment (The following analysis is not part of polymeval ‘reference’ mode). Therefore, we annotated putative non-B-DNA motifs in the *Carollia* reference genome with nonB-gfa [43], converting resulting tsv files to bed files and merging overlapping entries of the same annotation class with bedtools v2.30.0 [91]. Afterwards, we computed base coverage per breakpoint interval with bedtools coverage, using the breakpoint bed file obtained above as a reference. For each non-B-DNA annotation, we then plotted the mean number of bases annotated in that category (again using 200 bp non-overlapping windows) along aggregated breakpoint windows with a custom R script [63].

### Variant calling benchmarks for human samples

Apart from biodiversity genomics, ultra-low input sequencing can be a powerful approach for clinical samples such as biopsies or saliva, where the extractable HMW DNA is likewise limited [27]. Often, variant detection is of special interest in this context. Furthermore, while non-model organisms such as the ones we de-novo sequenced for polymerase benchmarking generally do not have curated, validated sets of Single-nucleotide variants (SNV), Insertion and Deletion (InDel), as well as structural variation (SV) or Tandem Repeat (TR) polymorphisms, these sets exist for the Genome in a Bottle (GIAB) human cell line, enabling us to test variant calling accuracy with a gold standard truth set. Following a recent study on human ULI sequencing [27], we therefore profiled the variant calling accuracy of amplified human reads (DeCodiFi, KODXtreme, repliQa. TaKaRaLongSeq). We furthermore added read sets sequenced from amplified human data previously [24,27] (Table S14).

To benchmark SNVs and small InDels, we aligned all reads (both full and downsampled to smallest coverage of the datasets sequenced in this study, KODXtreme, ∼25X) to the human reference genome hg38 with minimap2 v2.30 [45] and called variants with deepvariant v1.10.0 [46]. We then used <u>hap.py</u> v0.3.15 (https://github.com/Illumina/hap.py) with the truth set GIAB HG002 v4.2.1 and computed precision, recall and F1-score. Furthermore, to evaluate performance of larger structural variation, we aligned the same reads to the hg37 reference as before and ran sniffles v2.8.11 [48] with default parameters. We then used truvari v5.5.0 [92] to assess the same metrics as before against the HG002 Tier1 SV v0.6 benchmark set. For tandem repeat benchmarking, we called TR variants with trgt v5.1.0 [47], used the Tandem Repeat v1.0.1 ground truth variant set and computed precision, recall and F1-score with truvari v5.5.0, adding an additional ‘refine’ step. We additionally computed benchmarks against the CMRG v1.00 ground truth set [49] for small and structural variants, using the previously generated vcf files from deepvariant and additionally calling SVs against hg38 with sniffles as before, once more employing <u>hap.py</u> and truvari for precision and recall calculations.

To ensure robustness of the results above, we opted test one more, recently created ground truth set incorporating even more difficult regions and a more comprehensive structural variant set for hg38 (v5.0q, smallvar and SV, [50]), a recently published benchmarking tool, aardvark v1.0.0 [52], that allows differentiation between exact variant matches and basepair similarity (partial credit), as well as another variant calling tool, longcallD v0.0.11 [51], that can call small and structural variants at the same time. Consequently, we ran all possible combinations of variant caller, benchmarking engine, and ground truth set (Table S15-18, Figure 6). For a complete overview of which ground truth sets were used, refer to Table S26. The workflow is completely reproducible and integrated into polymeval with the ‘fetch_benchmarks’ (for downloading the ground truth files) and ‘variant_calling_benchmarks’ modes.

During polymerase amplification, certain haplotypes might be preferentially or uniquely amplified, potentially leading to a skewed allele representation (allelic imbalance) or complete allele dropout [87]. With whole-genome long reads and a relatively uniform coverage, we neglect potential overdispersion [93] and define an instance of allelic imbalance as a correctly called heterozygous variant whose alternative (reference) allele depth deviates significantly from the expectation of 0.5 x locus depth under a binomial distribution. Furthermore, we define allelic drop-out as any case of a true heterozygous variant not being represented by the correct reference and alternative alleles in a call set. This can include a variant not being called, several cases of tool-specific reference-(“RefCall”) or no-call (“NoCall”) judgments on a detected alternative variant, an incorrect alternative variant being called, or the alternative variant being called homozygous. Since the v5.0q smallvar ground truth set has the highest number of heterozygous variants [50], we used it as a reference for the following analysis. We normalized deepvariant vcf files of the downsampled reads (see above) with bcftools v1.24 [81], retaining only biallelic variants. Next, we tabulated the fate of each true heterozygous SNV and InDel from v5.0q in our read sets, using a custom R script [94] to assess the extent of allelic drop-out. Furthermore, we recorded the variant allele frequency of each correctly called variant, calculated as the number of reads assigned to the alternative allele by the caller (alternative allele depth) divided by the sum of reads assigned to both reference and alternative allele. We also used the alternative allele depth as the number of successes for a per-variant two-sided binomial test with p set to 0.5 and the number of trials set to overall variant depth, respectively. We then corrected for multiple testing using False discovery rate (FDR) correction. We note that at an average depth of 25X, the power to detect allelic imbalance is relatively low. We therefore also plotted VAF densities of SNVs, Insertions and Deletions and visually compared density curves of non-amplified and amplified reads (Figure 6D). The allelic imbalance analysis for deepvariant and longcallD called variants is implemented in the ‘variant_calling_benchmarks’ mode of polymeval.

To phase variants into haplotype blocks, we used HiPhase v1.7.0 [54], again with downsampled deepvariant-called vcf files (and the underlying mapped reads) as the input. We then recorded the number of variants that could be phased, as well as the overall haplotype block size N50 (Table S20).

